# diaPASEF Enables High-throughput Proteomic Analysis of Host Cell Proteins for Biopharmaceutical Process Development

**DOI:** 10.1101/2024.02.21.581388

**Authors:** Taku Tsukidate, Alyssa Q. Stiving, Shannon Rivera, Ansuman Sahoo, Sri Madabhushi, Xuanwen Li

**Author notes:** **Corresponding Author** T.T.; X.L.

## Abstract

Monitoring and quantifying host cell proteins (HCPs) in biotherapeutics production processes is crucial to ensure product quality, stability, and safety. Liquid chromatography–mass spectrometry (LC–MS) analysis has emerged as an important tool for identifying and quantifying individual HCPs. However, LC-MS–based approaches face challenges due to the wide dynamic range between HCPs and the therapeutic protein, as well as laborious sample preparation and long instrument time. To address these limitations, we evaluated the application of parallel accumulation–serial fragmentation combined with data-independent acquisition (diaPASEF), to HCP analysis for biopharmaceutical process development applications. We evaluated different library generation strategies and LC methods, demonstrating the suitability of these workflows for various HCP analysis needs such as in-depth characterization and high-throughput analysis of process intermediates. Remarkably, the diaPASEF approach enabled the quantification of hundreds of HCPs that were undetectable by a standard data-dependent acquisition mode while considerably improving sample requirement, throughput, coverage, quantitative precision, and data completeness.

---

Host cell proteins (HCPs) are protein impurities derived from host cells during the biotherapeutics production process. Several HCPs are considered high risk of affecting product quality, stability, and/or safety and require control strategies for removal and monitoring.^1–4^ The enzyme-linked immunosorbent assay (ELISA) employing anti-HCP polyclonal antibodies is the most used method to monitor total HCP amount during process development.^5^ However, since ELISA relies on immunoreactivity, it cannot detect non-immunogenic proteins or identify individual proteins. In this regard, liquid chromatography–mass spectrometry (LC–MS) analysis has become an important tool to identify and quantify individual HCPs.^6^

One substantial challenge encountered by LC-MS–based approaches is the wide (> 10^6^) dynamic range between the sparse HCPs and the abundant therapeutic protein in the drug substance. To enable LC-MS–based HCP analysis, we and others have reported various sample preparation and LC-MS strategies. For example, Protein-A affinity purification^7^ and molecular weight cutoff filtration^8^ deplete monoclonal antibodies (mAb) while chemical proteomics^9^ and ProteoMiner^10^ enrich certain HCPs. Alternatively, native digestion selectively digests proteolytically more labile HCPs over mAbs.^11^ In addition, offline fractionation^12,13^, long gradient LC^14^, and ion mobility spectrometry^15,16^ improve separation between HCP– and therapeutic protein–derived peptides. However, these strategies typically involve somewhat laborious sample preparation and/or require long instrument time (> 2 h / sample), which considerably limits sample throughput.^13^ Furthermore, datadependent acquisition (DDA), which remains to be the method of choice for HCP analysis, might not sufficiently cover the required dynamic range to allow for adequate detection or quantification of HCPs.^17,18^

Data-independent acquisition (DIA) is an attractive alternative to DDA.^19^ Typical DDA methods select the most abundant precursor ions of an MS1 scan for fragmentation and acquisition of MS2 spectra, leading to a loss of information, especially for sparse precursors, and partial data reproducibility due to the stochastic nature of precursor selection. In contrast, DIA methods slide pre-defined precursor isolation windows across the entire MS1 scan range for fragmentation and acquisition of MS2 spectra, enabling high reproducibility, sensitivity, and quantification across a broad dynamic range. Previous studies have applied various DIA methods to HCP analysis and demonstrated their advantages.^16,17,20–23^Among different DIA approaches, the diaPASEF method, which synchronizes ion mobility separation and mass selection to achieve great selectivity and sensitivity, is a transformative technology.^24,25^ However, the application of diaPASEF to HCP analysis has not yet been reported in peer-reviewed literature.

In this manuscript, we report the first application of diaPASEF to HCP analysis in the biopharmaceutical industry and describe our optimized workflows (**Figure 1**). We systematically evaluated several spectral library generation strategies and LC methods using our internal pipeline molecules and identified workflows suitable for various HCP analysis needs in biotherapeutics process research and development.

**Figure 1.**
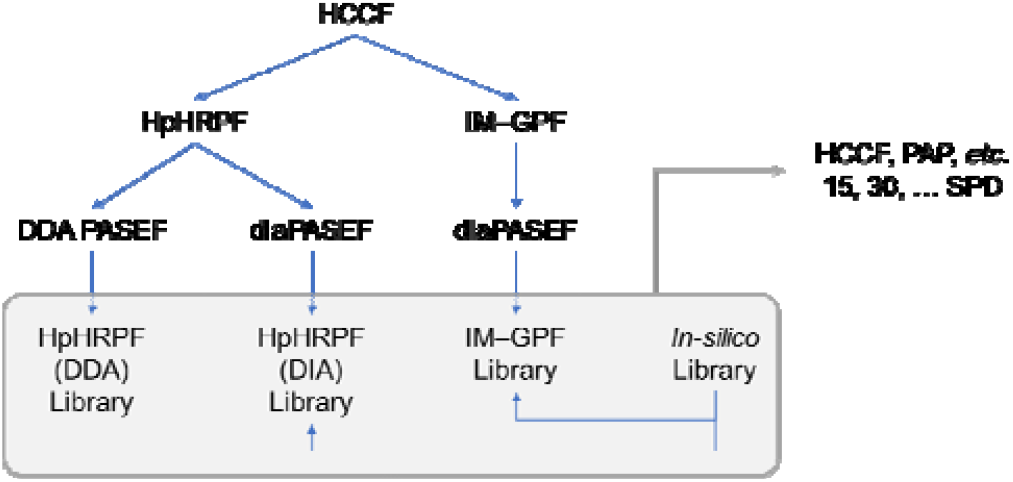
Overview of diaPASEF–based HCP analysis workflows. Spectral libraries were generated from HCCF employing different fractionation strategies and acquisition modes or *in silico*. These spectral libraries were used to process diaPASEF data of biotherapeutics process samples that were acquired with several LC methods. HCCF: harvested cell culture fluid; HpHRPF: high pH reversed-phase fractionation; IM–GPF: ion mobility–coupled gas-phase fractionation; PAP: protein A pool; SPD: samples per day.

## Materials and Methods

### Protein samples

Pierce Hela Protein Digest Standard was purchased from Thermo Scientific. mAb1 HCCF, mAb2 HCCF, and mAb1 PAP were internally sourced.

### Proteomics sample preparation from mAb1 or mAb2 HCCF

mAb1 or mAb2 HCCF (20 μL) was incubated with 4 M urea, 25 mM TCEP-HCl, and 50 mM acrylamide in 50 mM tris-HCl pH 8.0 (total volume: 50 μL) at 60 °C for 30 min. An aliquot (20 μL) was incubated with Sera-Mag SpeedBead (10 μL, 200 μg) and acetonitrile (70 μL) at 20 °C for 10 min with agitation at 500 rpm. The beads were washed three times with 80 % EtOH (200 μL) and then incubated with 50 mM tris-HCl pH 8.0 (20 μL) containing 0.4 μg trypsin and 0.1 μg LysC at 37 °C for 18 h. Peptide concentration was determined with the Nanodrop A280 assay and/or the BCA assay (Thermo Scientific) and adjusted to 0.02 g/L with 0.1 % formic acid before loading 20 μL (400 ng) on Evotip Pure Evotips (Evosep) according to the manufacturer’s instructions. The remainder was fractionated with Pierce High pH Reversed-Phase Peptide Fractionation Kit (Thermo Fisher Scientific), concentrated to dryness, reconstituted with 0.1 % formic acid to the final concentration of 0.02 g/L, and loaded (400 ng) on Evotip Pure Evotips according to the manufacturer’s instructions.

### Proteomics sample preparation from mAb1 PAP

mAb1 PAP were subjected to native digestion as previously described^11^. Peptide concentration was adjusted to 0.02 g/L (based on the initial input amount) with 0.1 % formic acid before loading 20 μL (400 ng) on Evotip Pure Evotips according to the manufacturer’s instructions.

### LC–MS analysis

Peptides were separated on the Evosep One LC system with Pepsep C18 columns using the manufacturer’s pre-defined methods denoted by daily sample throughput e.g., 30 samples per day (SPD) and analysed on the Bruker timsTOF Pro 2 system. The standard DDA PASEF method was used without modification. The diaPASEF method was optimized by adjusting the isolation windows either manually or with the py_diAID software^26^. The IM–GPF schemes were designed with a custom R script.

### Data analysis

DDA PASEF data were processed with MaxQuant v2.4.10^27^. Use of .NET Core was disabled. Match between runs (MBR) was disabled for spectral library generation and enabled for all other analyses. All other settings were left as default. MaxQuant evidence and msms files were reformatted with a custom R script and used to generate a spectral library with DIA-NN v1.8.1^28^. All diaPASEF data were processed with DIA-NN v1.8.1. Mass accuracies were set to 10 ppm for spectral library generation and to 15 ppm for all other processing. MBR was disabled for spectral library generation and enabled for all other analyses as MBR tended to have a small but positive effect even when processing data with IM-GPF or HpHRPF libraries (**Supporting Information Figures S1 & S2**). For the generation of the *in silico* spectral library from sequence databases, the precursor charge range and the mass range were restricted to 2−3 and 300–1200 Th, respectively. The global precursor and global protein FDR thresholds were set to 1 % for all analyses. This was achieved by filtering DIA-NN main reports on the Global.Q.Value column and the Global.PG.Q.Value column in MBR-disabled data processing and on the Lib.Q.Value column and the Lib.PG.Q.Value in MBR-enabled data processing. FDR estimation was independently validated by the two-species library analysis as previously described^29^ (**Supporting Information Figure S3**). The data post-processing analysis was performed in R version 4.0.3. Protein quantification was performed using the MaxLFQ algorithm as implemented in the iq package^30^ based on the Fragment.Quant.Corrected column of DIA-NN main reports. Chromatograms were visualized in Skyline-daily v19.0.9.

## Results and Discussion

### Optimization of ion mobility–coupled gas phase fractionation scheme

Choice of spectral library has significant impact on DIA MS data quality as well as total experiment time.^31–33^ For example, Wen et al. reported that a spectral library generated via HpHRPF and DDA yielded 10–20 % more identifications than an *in silico* spectral library in their diaPASEF analysis of human and mouse cell lysates.^32^ On the other hand, HpHRPF results in sample matrices that are different from the target samples, which may affect retention time accuracy. In this regard, gas-phase fractionation (GPF) is an attractive alternative to rapidly generate spectral libraries.^33^ For example, Penny et al. demonstrated that the ion mobility–coupled GPF (IM–GPF) workflow performed comparably with the HpHRPF workflow in terms of protein identifications in the HeLa cell protein digest.^34^ Similarly, Rice and Belani demonstrated the IM–GPF workflow outperformed the HpHRPF workflow in terms of protein identifications in depleted plasma samples.^35^ While both studies employed the isolation window width of 5 Th for their GPF library generation, narrower windows are desirable to improve the selectivity of precursor selection. Thus, we optimized the isolation window width and the number of PASEF scans for the IM–GPF scheme with the commercial HeLa protein digest standard (**Figure 2**).

**Figure 2.**
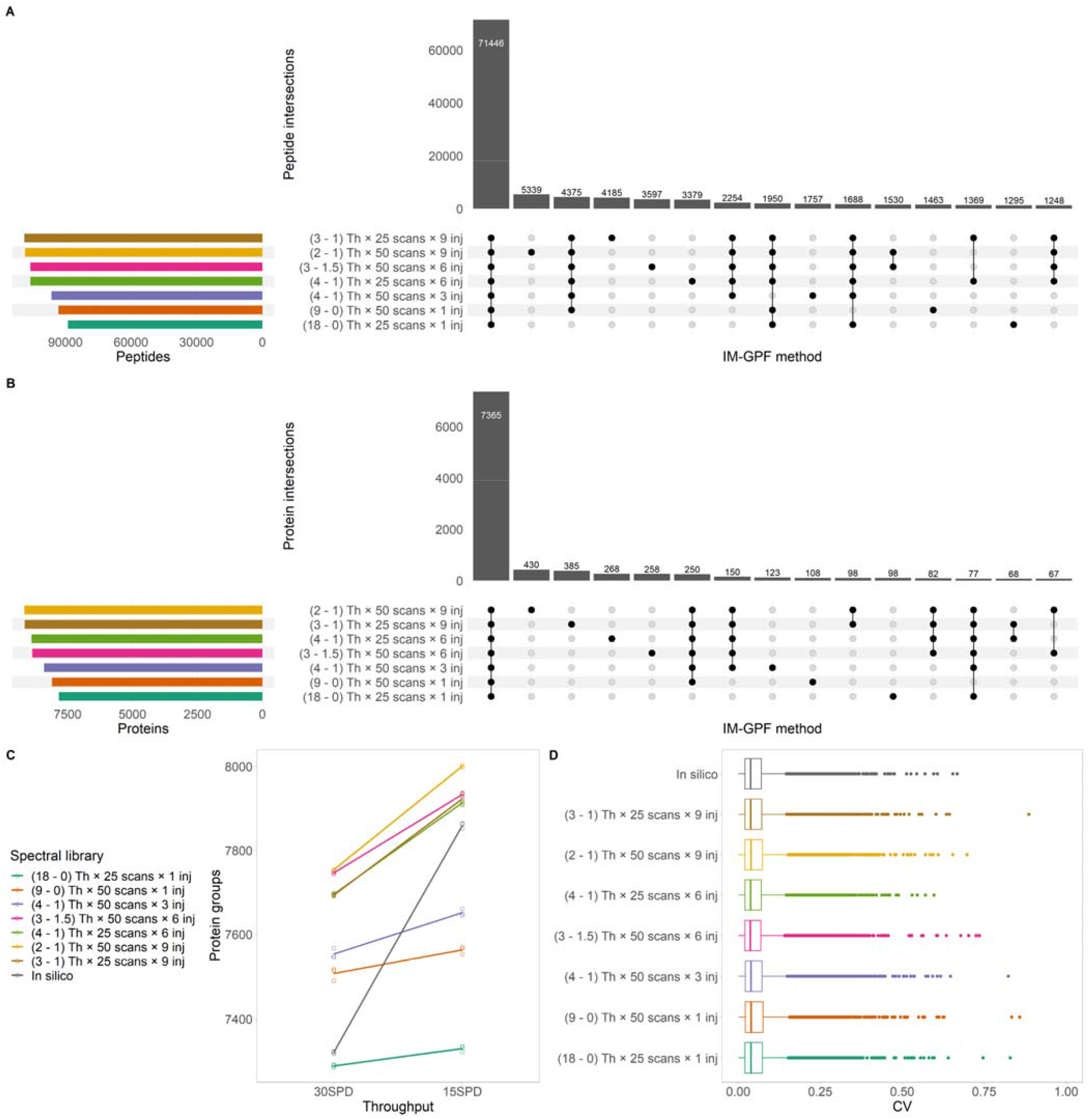
Optimisation of IM–GPF scheme with HeLa protein digest standard. The precursor isolation window width and the number of PASEF scans were varied. Number and overlap of peptides (A) and proteins (B) in various IM–GPF libraries. Horizontal bars indicate the number of peptides or proteins in each library. Vertical bars indicate the number of peptides or proteins shared by several libraries (intersection) as indicated by dotes below. Fifteen largest intersections are shown. Number of quantified protein groups (C) and coefficients of variation from the 30 SPD data (D) in three replicate injections, employing the IM–GPF and *in silico* libraries during data processing. The IM–GPF scheme names and corresponding library names represent (window width – overlap) Th **×** (number of PASEF scans) **×** [number of required injections (abbreviated as inj)]. Evosep method: 15 SPD for spectral library data acquisition; 15 or 30 SPD for three replicate injections. Global peptide and global protein FDRs < 0.01.

The IM–GPF schemes with narrower isolation windows yielded larger spectral libraries with more peptide and protein identifications but required more injections to cover the whole mass range (**Figure 2A–B**). The methods with 50 PASEF scans yielded 2.5–3 data points per peak at full width at half maximum (FWHM) on average whereas the methods with 25 PASEF scans yielded ∼ 4 data points per peak at FWHM on average, suggesting that these schemes may also be suitable for shorter gradient LC methods (*vide infra*).^36^ As expected, the libraries shared most of the library peptides and proteins. On the other hand, the use of different libraries for the analysis of HeLa diaPASEF data resulted in modest difference in the numbers of protein identifications (**Figure 2C**). The larger libraries (6 or 9 injections) consistently outperformed the smaller libraries (1 or 3 injections) and an *in-silico* library generated by DIA-NN in terms of protein identifications. Protein quantification was highly precise with median coefficients of variation of 3–4 % and unaffected by the choice of spectral library (**Figure 2D**). Interestingly, the “(3 – 1.5) Th **×** 50 scans **×** 6 injections” library outperformed the “(3 – 1) Th **×** 25 scans **×** 9 injections” library, highlighting the importance of specificity in spectral library^37^. Overall, applying the effective 1-Th isolation windows as in the “(2 – 1) Th **×** 50 scans **×** 9 injections” scheme resulted in the best performance (**Supporting Information Figure S4**).

### Optimisation of spectral library workflow for biotherapeutics process intermediates

With the optimized IM–GPF scheme in hand, we evaluated several spectral library generation workflows with a harvested cell culture fluid (HCCF) from one of our proprietary monoclonal antibody processes (mAb1, **Figure 3, Supporting Information Figure S5**). We processed mAb1 HCCF with one-step denaturation, reduction and alkylation followed by the SP3 cleanup and on-bead digestion to prepare a protein digest.^38^ We then subjected the digest to the offline HpHRPF employing a commercial kit, which generates 10 fractions, or to the optimized IM–GPF scheme to generate spectral libraries. The HpHRP fractions were analysed with a standard DDA PASEF method (HpHRPF–DDA) or a diaPASEF method with non-overlapping 9-Th isolation windows and 50 PASEF scans (HpHRPF–DIA). The cycle time and hence the number of data points per peak of this diaPASEF method matches those of the optimized IM–GPF method under the same LC gradient. The HpHRPF–DIA workflow resulted in the largest spectral library including > 16,000 peptides (**Figure 3A**) and > 1,800 proteins (**Figure 3B**) that are unique to this library and not part of the IM–GPF or HpHRPF–DDA libraries. On the other hand, the HpHRPF–DDA workflow resulted in the smallest spectral library. Accordingly, the use of the HpHRPF–DIA library for HCCF diaPASEF data processing resulted in the largest number of protein identifications (**Figure 3C**). In contrast to the HeLa diaPASEF data, the IM–GPF libraries underperformed an *in-silico* library generated by DIA-NN in terms of protein identifications in the HCCF diaPASEF data acquired with the 15 SPD method. However, the IM-GPF library advantageously outperformed the *in-silico* library in the HCCF diaPASEF data when increasing sample throughput. For example, processing 100SPD diaPASEF data with either the IM-GPF (15 SPD) library or the HpHRPF (15 SPD, DIA) library yielded similar number of protein identifications and abundance distributions (**Figure 3D**). This is likely because many of those peptides that are unique to the HpHRPF (15 SPD, DIA) library were not resolved well enough with the short gradient from more abundant peptides that are also part of the IM-GPF (15 SPD, DIA) library. On the other hand, protein abundance distribution clearly shifted towards more abundant proteins when the same data were processed with the HpHRPF (15 SPD, DDA) library, perhaps reflecting potential abundance bias in the DDA-based library. Quantitative precision was largely unaffected by the choice of spectral library (median CV ∼ 6 %).

**Figure 3.**
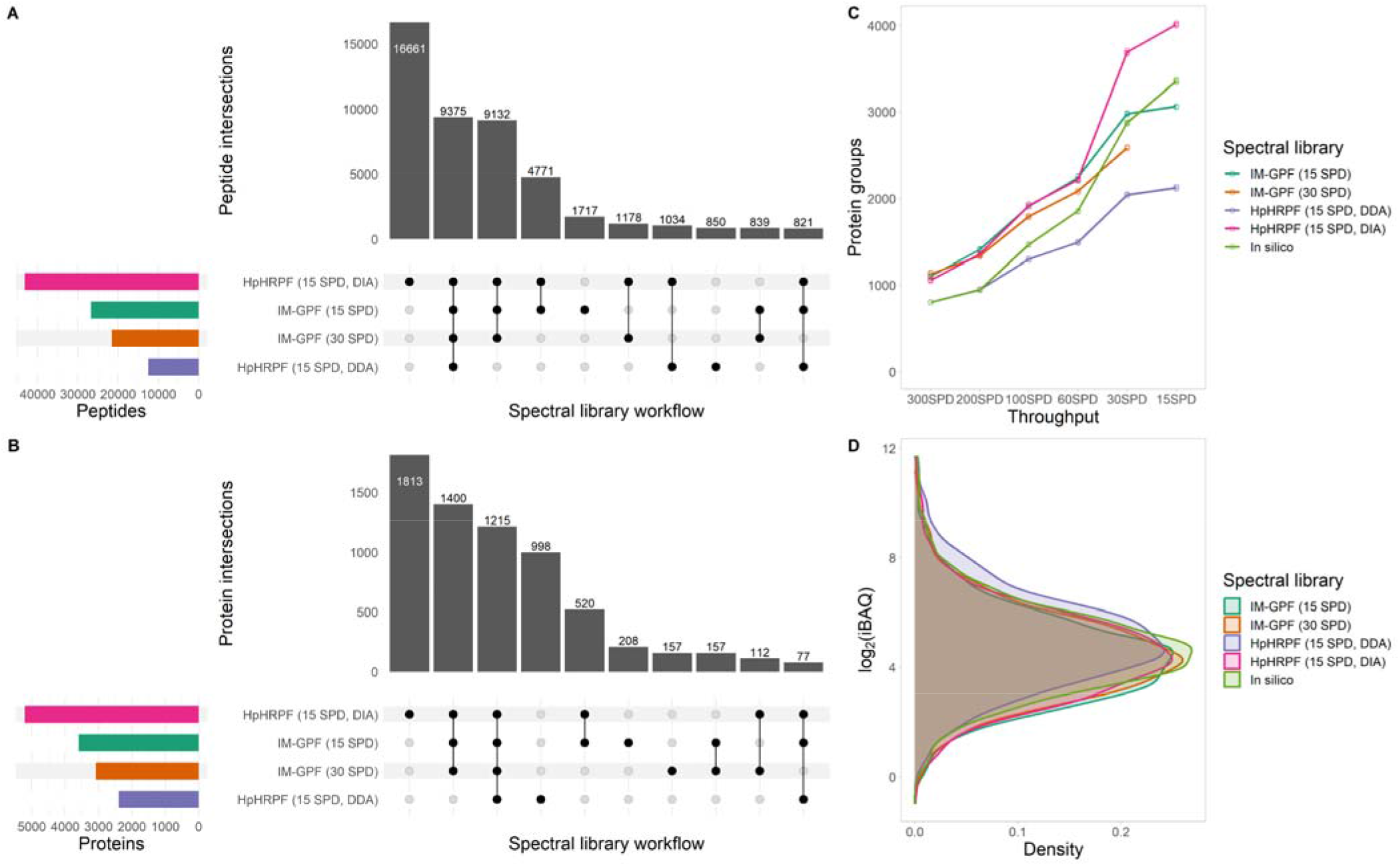
Evaluation of spectral library generation workflows with mAb1 HCCF. Empirical spectral libraries were generated by the optimal IM–GPF scheme or HpHRPF combined with DDA PASEF or diaPASEF. Number and overlap of peptides (A) and proteins (B) in various libraries. Horizontal bars indicate the number of peptides or proteins in each library. Vertical bars indicate the number of peptides or proteins shared by several libraries (intersection) as indicated by dotes below. Ten largest intersections are shown. Number of quantified protein groups in three replicate injections (C) and distribution of average protein abundance score (iBAQ) (D), employing the empirical and *in-silico* libraries during data processing. D: 100 SPD. Global peptide and global protein FDRs < 0.01.

Similarly, we evaluated the performance of these spectral libraries for the analysis of mAb1 protein A pool (PAP) diaPASEF data (**Figure 4, Supporting Information Figures S6 & S7**). We processed mAb1 PAP according to a native digestion protocol as previously described^11^. The HpHRPF– DIA and the IM–GPF (15 SPD) libraries led to similar protein identifications (**Figures 4A**) and quantitative precisions (median CV ∼ 12 %). On the other hand, the use of HpHRPF– DDA (median CV ∼ 14 %) or the *in-silico* libraries (median CV ∼ 10 %) resulted in significantly smaller numbers of protein identifications. Of note, protein abundance distribution clearly shifted towards more abundant proteins in the case of the *in-silico* library (**Figure 4B**), highlighting the importance of experimental spectral library in the diaPASEF analysis of biotherapeutics process samples to avoid the possibility of missing lower abundance HCPs.

**Figure 4.**
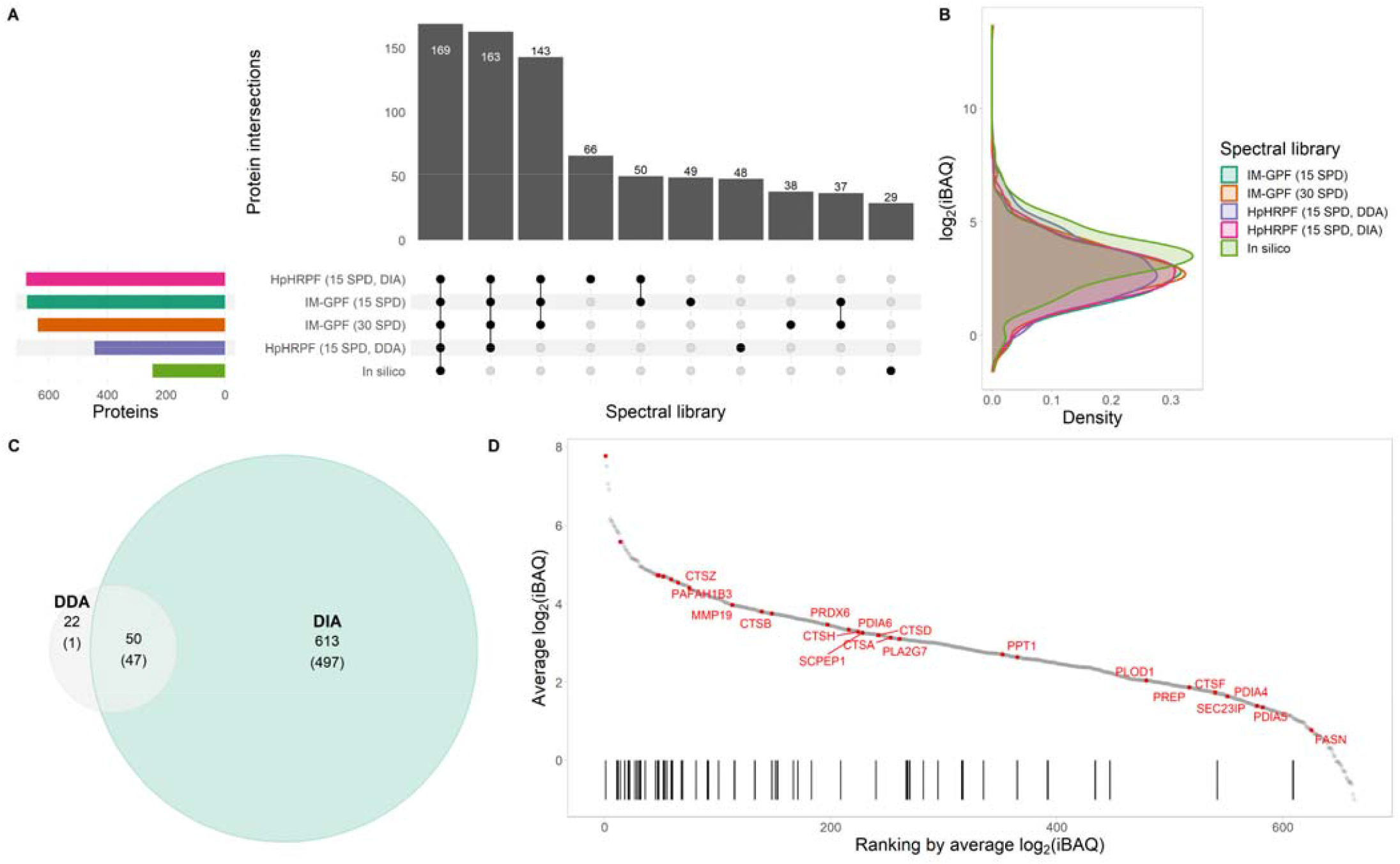
Evaluation of spectral library generation workflows with mAb1 PAP. Number and overlap of protein groups that were quantified in all three replicate injections (A) and distribution of average protein abundance score (iBAQ) (B), employing the empirical HCCF and *in silico* libraries during data processing. In A, horizontal bars indicate the number of proteins; vertical bars indicate the number of shared proteins (intersection); ten largest intersections are shown. (C) Comparison of HCP identifications between DDA PASEF and diaPASEF methods. The numbers in parentheses indicate the numbers of proteins with at least two unique peptides. (D) Waterfall plot of average protein abundance score for the 663 HCPs quantified with the diaPASEF method. High-risk HCPs are highlighted as red points and, if not quantified with the DDA PASEF method, are labelled with their protein symbols. The black bars indicate quantification of the HCP with the DDA PASEF method and the DDA method’s significant (p = 2 **×** 10^−8^, Kolmogorov-Smirnov test) bias towards abundant HCPs. 30 SPD. Global peptide and global protein FDRs < 0.01.

Next, we compared the diaPASEF method and the standard DDA PASEF method in the analysis of the mAb1 PAP sample. The diaPASEF method quantified a majority (50 of 72) of proteins quantified by the DDA PASEF method and 613 additional HCPs (**Figure 4C**). The 22 proteins that were quantified by the DDA PASEF method but not by the diaPASEF were supported only by 24 peptides in total, of which 23 peptides were not part of any of the four experimental spectral libraries. Visual inspection of corresponding spectra revealed many questionable assignments. Moreover, none of these 22 proteins are considered high-risk HCPs.^1^ On the other hand, diaPASEF revealed the presence of > 600 additional HCPs that the DDA PASEF method missed (**Figure 4D**). These HCPs included several high-risk HCPs and polysorbate-degradative enzymes such as PLA2G7 and PPT1. In addition, quantification was more precise with the diaPASEF method (median CV ∼ 12 %) than with the DDA PASEF method (median CV ∼ 16 %).

Furthermore, the diaPASEF method was clearly superior to our platform method that employs a three-hour gradient on a microflow LC and leverages conventional DDA on the Q Exactive HF-X mass spectrometer^2^—The new method required 100-fold less sample, increased throughput by > 10-fold, expanded identification coverage by up to 10-fold, considerably improved quantitative precision, and improved data completeness by five-fold (**Supporting Information Figure S8**).

GO term analysis of mAb1 HCCF and mAb1 PAP suggested that most HCP mass (∼ 90 %) was released from plasma membrane, cytoplasm and/or nucleus through cell rapture and that (actively secreted) extracellular proteins accounted for a small fraction of the total HCP mass (∼ 10 %) (**Supporting Information Figure S9**). In terms of HCP persistence, the abundance in HCCF and the molecular weight of HCP were strongly correlated with the abundance of HCP in PAP: ρ_HCCF_ = 0.79; ρ_MW_ = −0.81. On the other hand, the isoelectric point and the hydrophobicity index lacked correlation with the abundance in PAP. In addition, distributions of pI or GRAVY values were similar between HCCF and PAP. These results extend similar findings in previous studies beyond the limited numbers of HCPs to many other diverse HCP populations.^39,40^ In addition, the HCP peptides overall eluted earlier than the HeLa protein digest, which may help LC method optimization. Altogether, these results indicate that the diaPASEF method combined with rapid IM–GPF spectral library generation enables wide proteome coverage in biotherapeutics process samples.

### High-throughput diaPASEF analysis of biotherapeutics production process intermediates

To assess the potential of the optimal diaPASEF workflow to further advance biotherapeutics process understanding, we tested its performance in a high-throughput analysis of mAb2 HCCFs (**Figure 5**). These HCCF samples originated from several mAb2 clones that were cultured under various conditions and harvested at several time points, covering 72 different conditions. We generated a spectral library from 18 repeated IM-GPF measurements of a sample pool with the effective 1-Th isolation window *i*.*e*., “(2 – 1) Th **×** 25 scans **×** 18 injections” method to balance the optimal isolation window width and cycle time and analysed the 72 single-shot samples in addition to 12 blanks, 12 system suitability samples, and 11 sample pools in a random order at the 100-SPD throughput. Despite the large number of runs required to accomplish this experiment, with the 60 and 100 SPD methods this took < 36 hours from start to finish. Sample pools were highly reproducible with the correlation coefficient (Pearson’s r) > 0.97 between any two runs and median CV ∼ 6 %, demonstrating the excellent performance of the method. Sample carry-over effect was virtually absent. We quantified ∼ 1800 HCPs with few missing values (**Figure 5A**) including ∼ 30 high-risk HCPs^1^ such as PLA2G7 (**Figure 5B**). Culture condition, time point, and clone had clear impact on overall HCP profiles and high-risk HCP levels. For example, PLA2G7 was more abundant in the HCCFs of cells cultured under the condition A than in the HCCFs of cells cultured under the condition B. In addition, we identified a cluster of HCPs to which most high-risk HCPs belong. Further analysis of this data will be presented elsewhere (Sahoo *et al*., manuscript in preparation).

**Figure 5.**
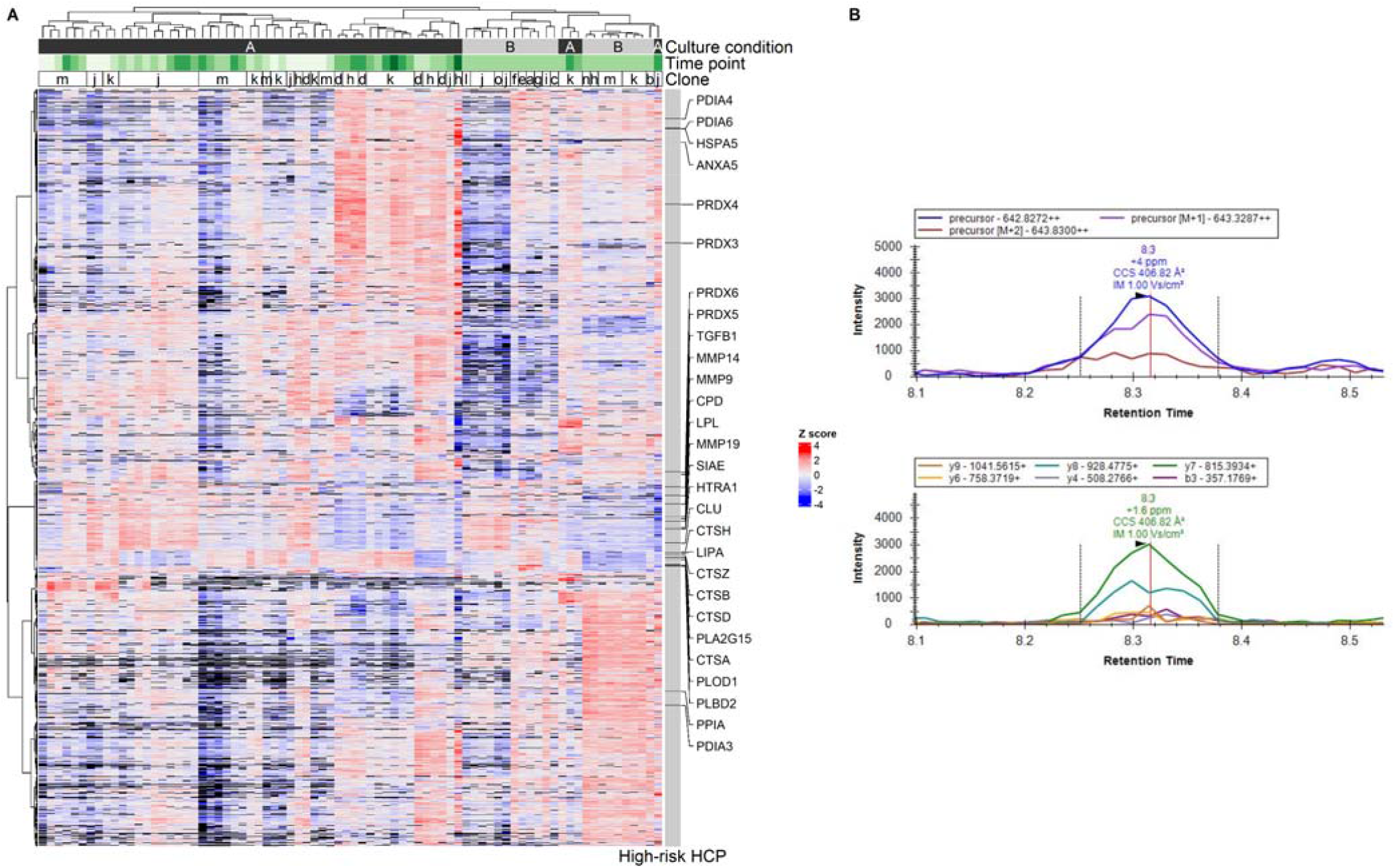
High-throughput diaPASEF analysis of mAb2 HCCFs. (A) Heatmap visualization of protein quantity. Each row corresponds to a protein group and each column corresponds to a sample. Fill colour indicates row-wise Z score. High-risk HCPs are indicated with black bars in the annotation panel on the right. Sample pools were highly reproducible with the correlation coefficient (Pearson’s r) > 0.97 between any two runs and median CV ∼ 6 %. (B) A representative extracted ion chromatogram for the PLA2G7-derived peptide ENILGSYFDVK and its MS2 fragments. Global peptide and global protein FDRs < 0.01.

*In-silico* libraries could be advantageous when analyzing a large set of heterogenous samples. Indeed, we observed modest (< 10 %) increase in the number of quantified HCPs compared to the IM-GPF approach (**Supporting Information Figure S10**). However, data processing time increased considerably from 134 min for the IM-GPF approach including the time for processing the 18 IM-GPF files to 2082 min for the library-free approach. Thus, both approaches are valid options depending on study goals.

## Conclusions

In summary, we optimized a diaPASEF-based workflow for HCP analysis and demonstrated its potential to advance biotherapeutics process understanding. We evaluated several spectral library generation strategies and LC methods for biotherapeutics process samples and identified workflows suitable for various HCP analysis needs including high-throughput analysis of harvests and in-depth characterization of purification process intermediates. With an optimized workflow in hand, we demonstrated that the diaPASEF method can reveal the presence of hundreds of HCPs in a PAP sample that are undetectable by the standard DDA PASEF method including several high-risk HCPs and should facilitate investigative studies. We also demonstrated high-throughput analysis of HCCF samples, which should advance our understanding of HCPs prior to purification and inspire new control strategies.

## ASSOCIATED CONTENT

### Supporting Information

The Supporting Information is available free of charge on the ACS Publications website.

Additional experimental details and results (PDF)

## AUTHOR INFORMATION

### Author Contributions

The manuscript was written through contributions of all authors. All authors have given approval to the final version of the manuscript.

## ACKNOWLEDGMENT

T.T. and A.S. thank MRL Postdoctoral Program. The authors thank Douglas D. Richardson and Hillary A. Schuessler for guidance and support.

---

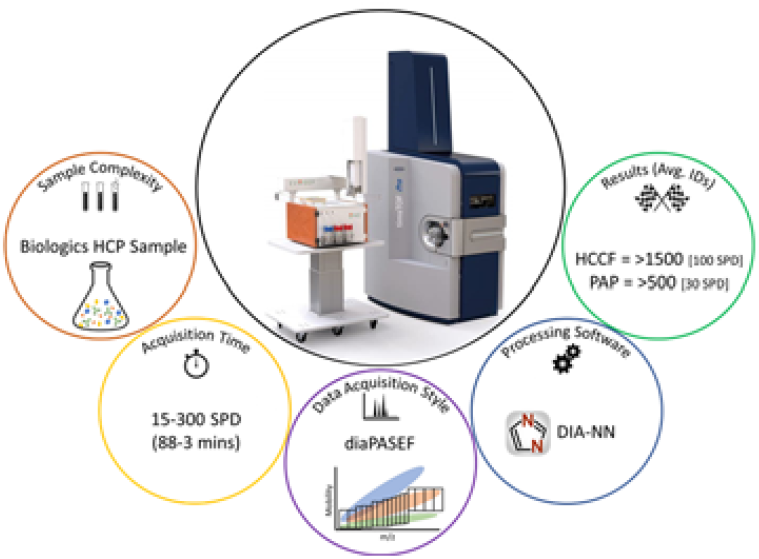
For Table of Contents Only

