## Supporting Information for "diaPASEF Enables High-throughput Proteomic Analysis of Host Cell Proteins for Biopharmaceutical Process Development"

**Table of Content**

Figures S1

Figures S2

Figures S3

Figures S4

Figures S5

Figures S6

Figures S7

Figures S8

Figures S9

Figures S10

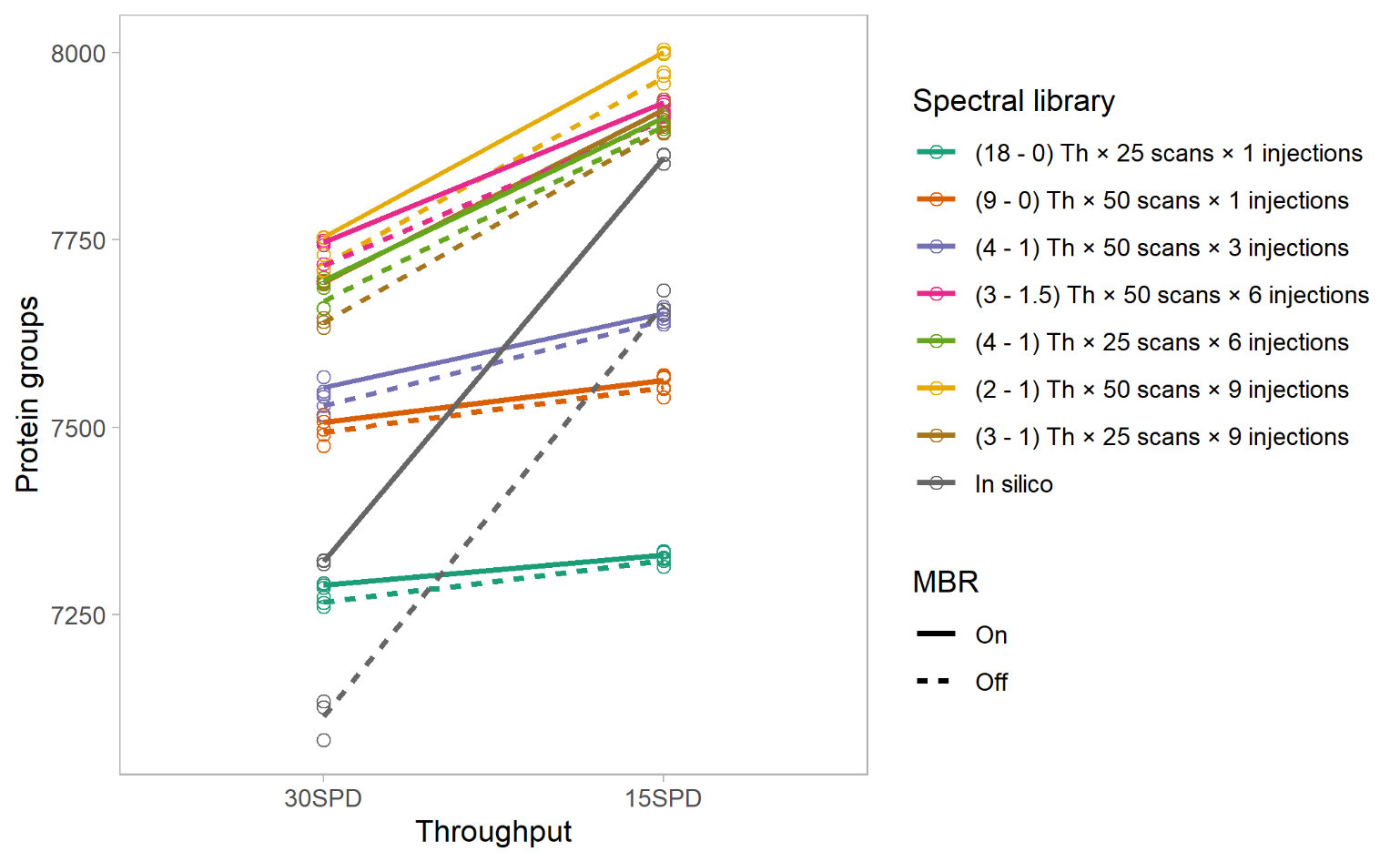

**Figure S1.** Impact of MBR on protein identification numbers in HeLa protein digest standard data processing with IM-GPF or *in-silico* libraries.

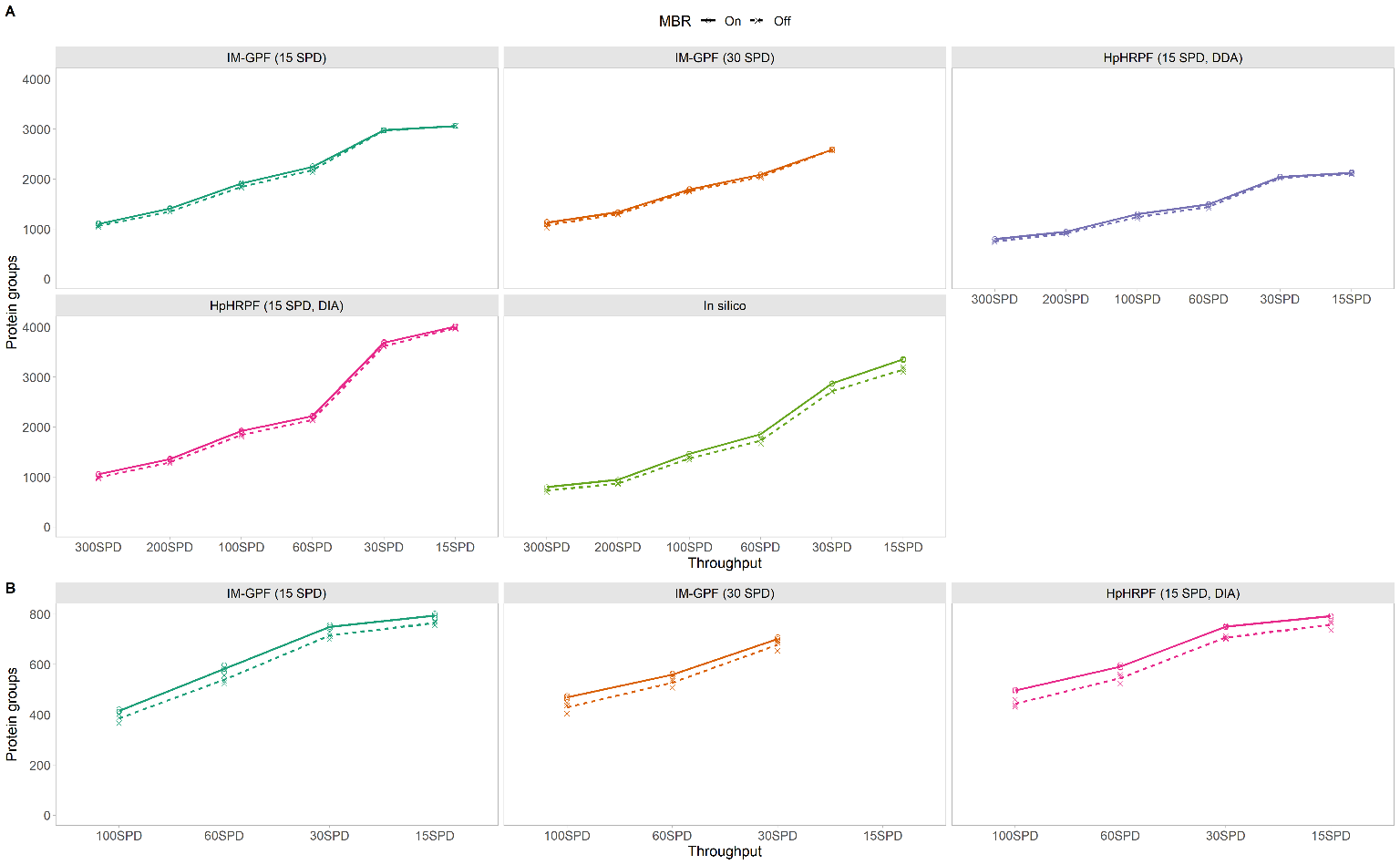

**Figure S2.** Impact of MBR on protein identification numbers in mAb1 HCCF (A) and mAb1 PAP (B) data processing with IM-GPF, HpHRPF, or *in-silico* libraries.

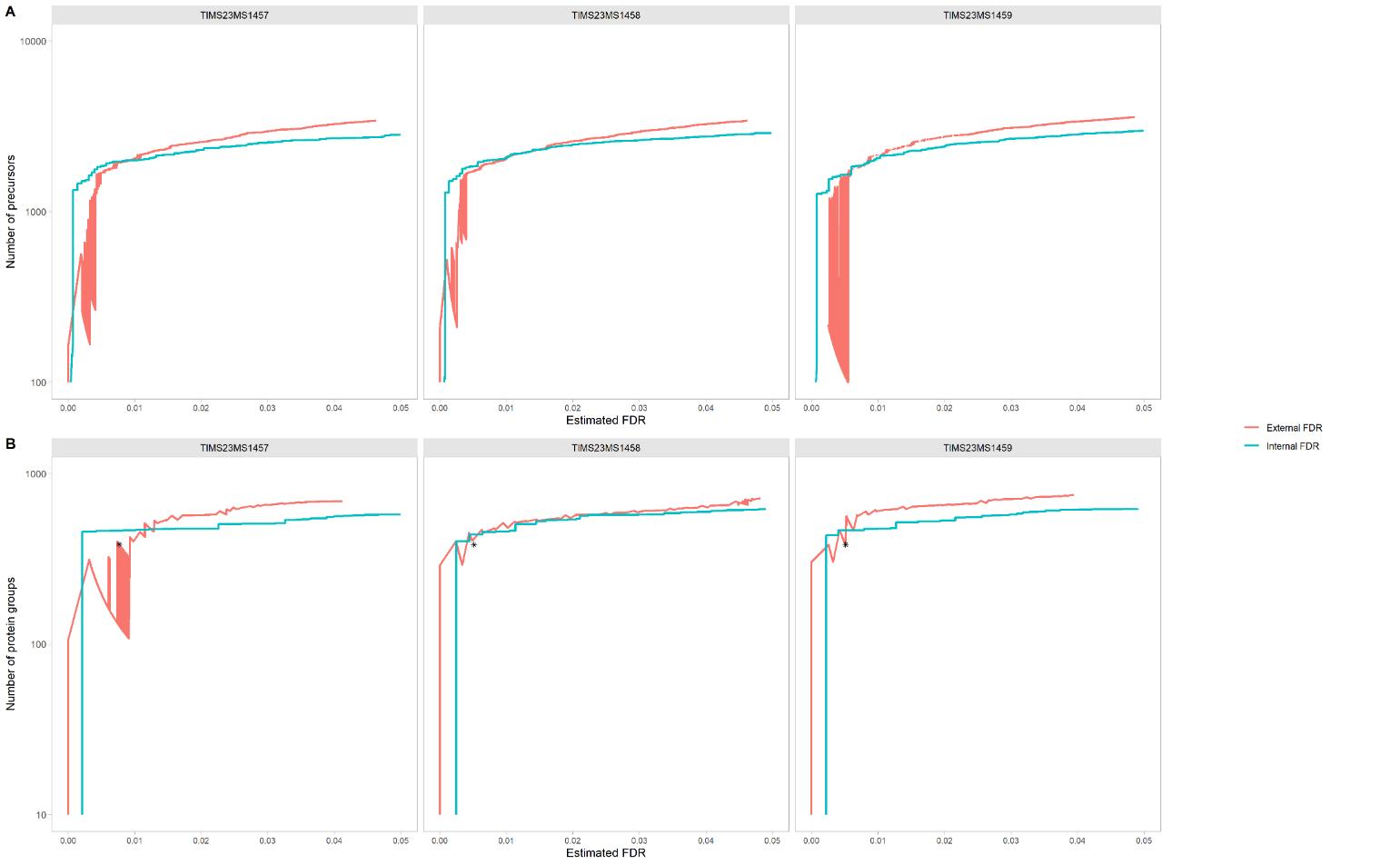

**Figure S3.** Validation of FDR estimation. External FDR and internal FDR showed good agreement at both precursor (A) and protein (B) levels. The asterisk indicates the external FDR and the number of protein groups after filtering the data by requiring precursor FDR < 0.01 and unique peptides ≥ 2 (“two-peptide rule”). The mAb1 PAP data were processed with a HpHRPF library generated from mAb1 HCCF and a yeast protein digest.

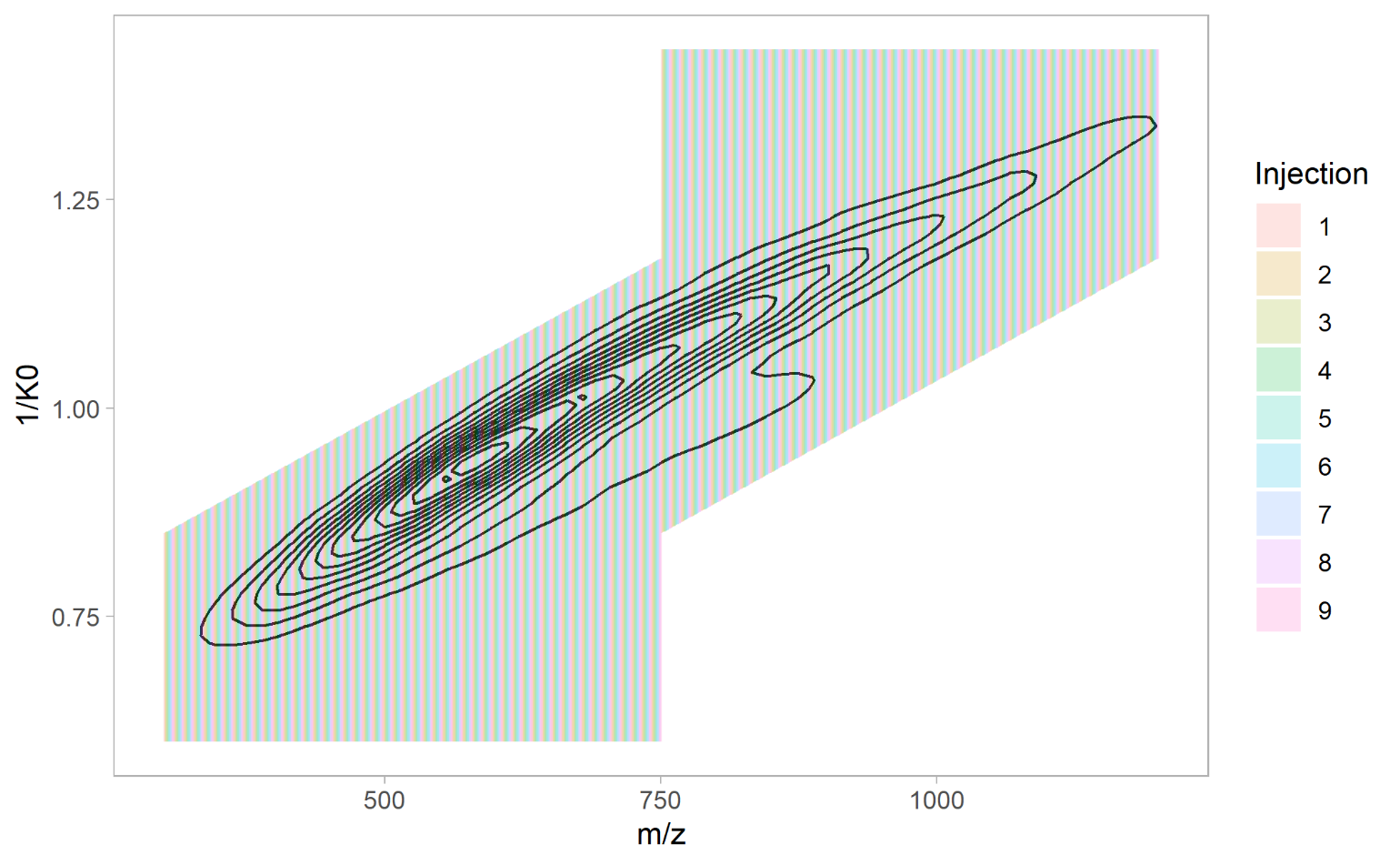
**Figure S4.** The IM–GPF scheme (2 – 1) Th × 50 scans × 9 injections overlaid on charges 2 and 3 precursor density in the HeLa protein digest.

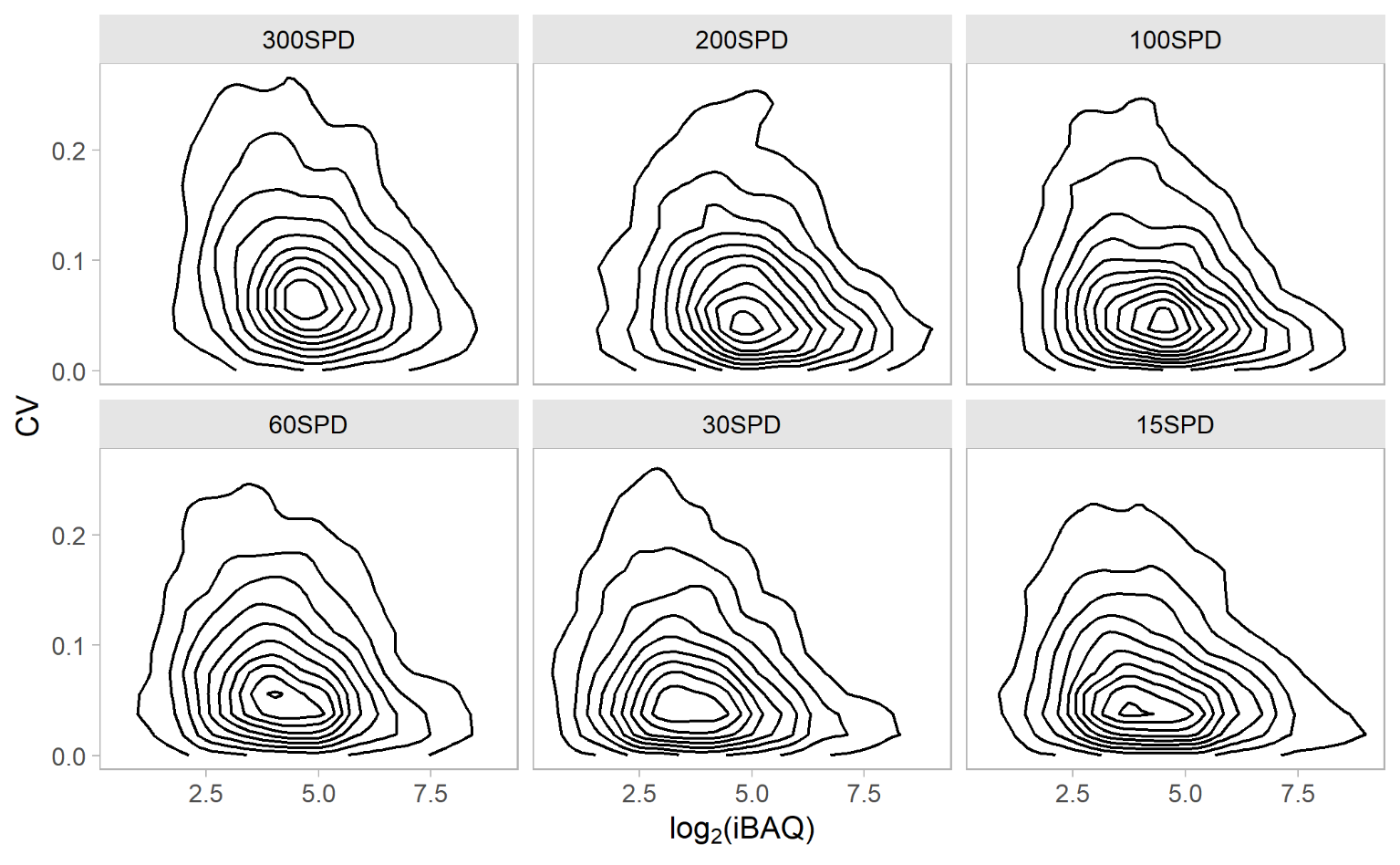

**Figure S5.** Relationship between protein abundance and coefficient of variance. Mean iBAQ scores and CVs were calculated based on three replicates of single-shot analysis of mAb1 HCCF. IM–GPF (15 SPD) library.

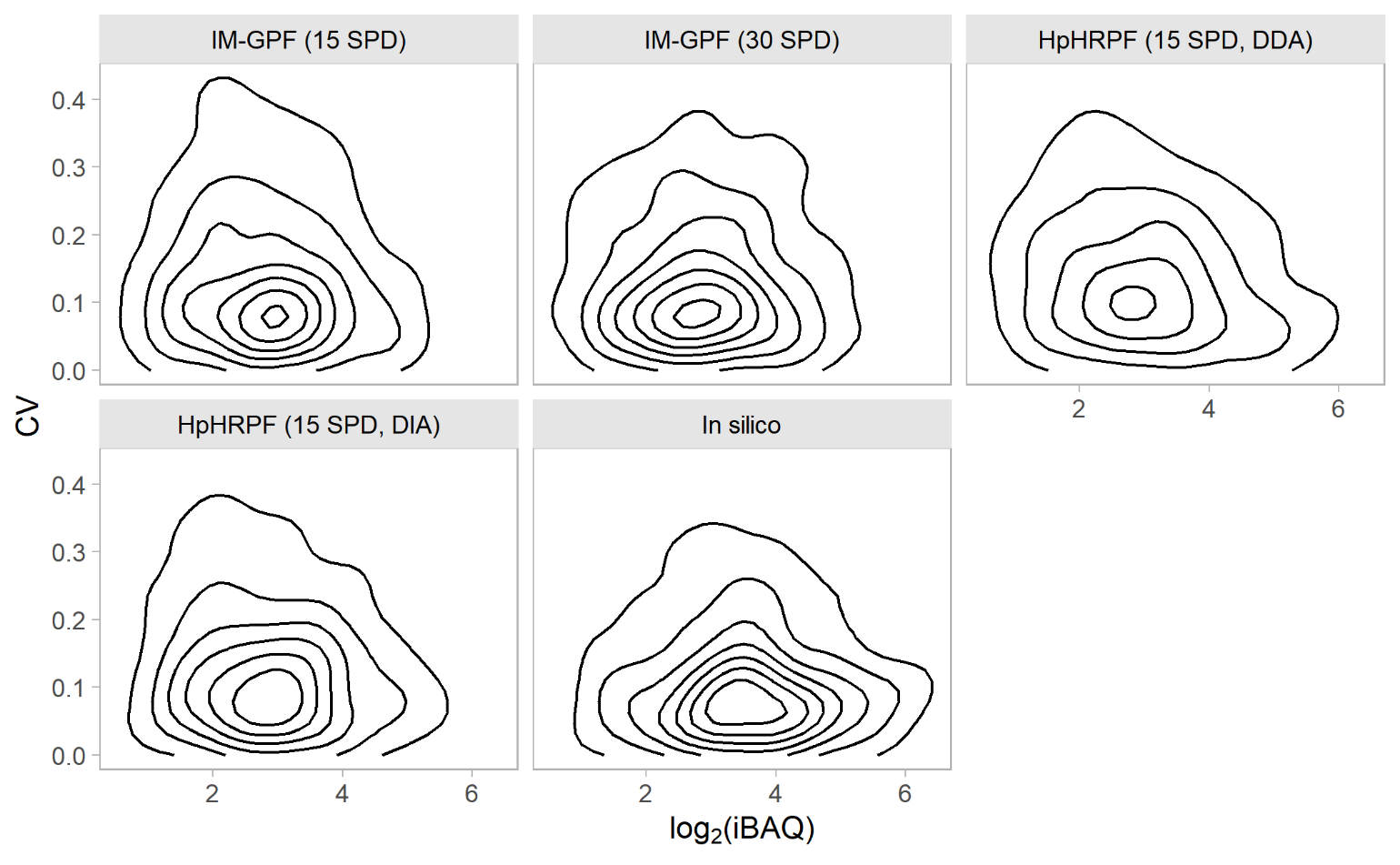

**Figure S6.** Relationship between protein abundance and coefficient of variance. Mean iBAQ scores and CVs were calculated based on three replicates of single-shot analysis of mAb1 PAP (30SPD).

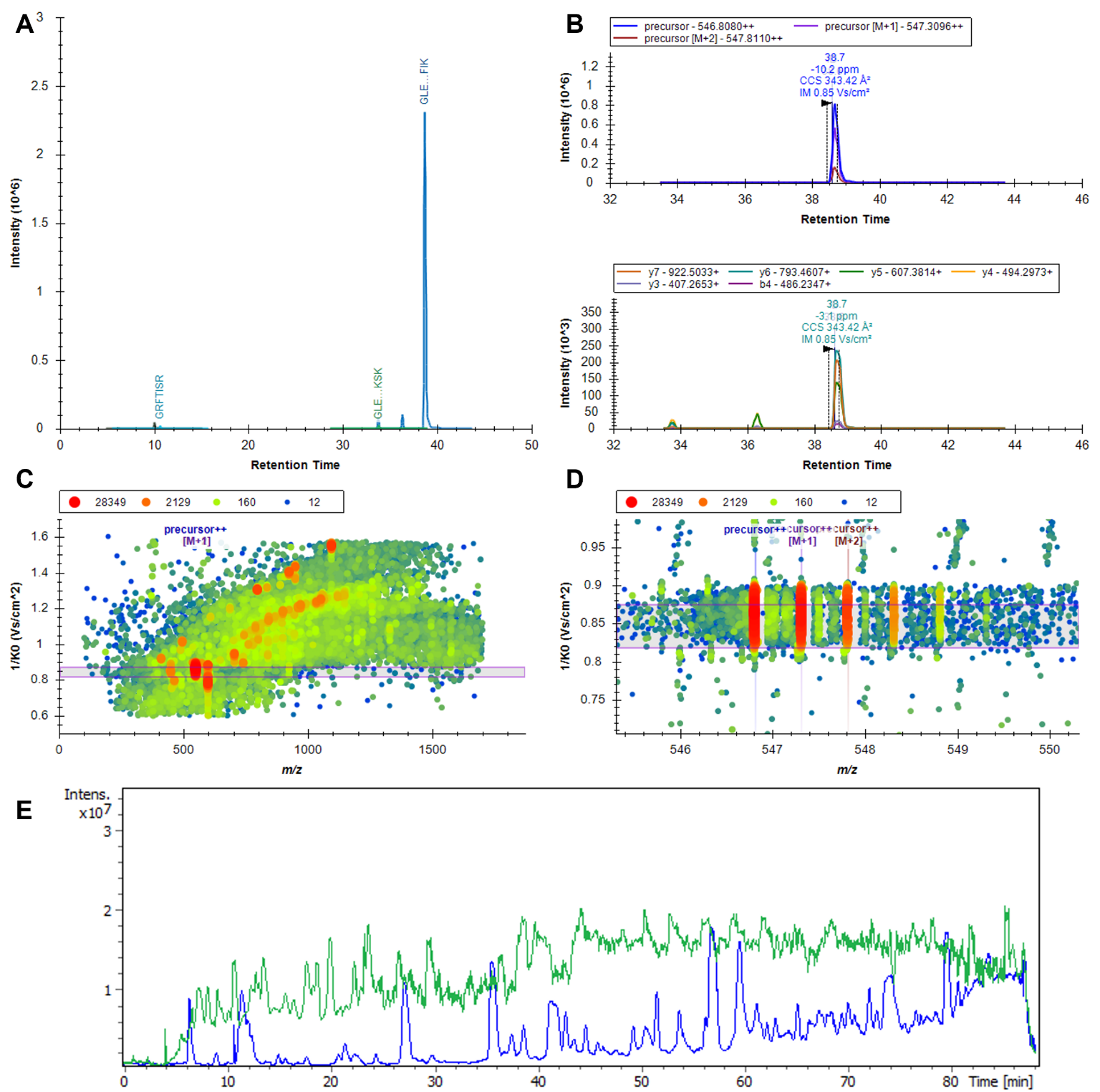

**Figure S7.** Example chromatogram of a mAb1-derived peptide. (A) Retention time and intensity of mAb1 heavy chain–derived peptides identified in mAb1 PAP. (B) Precursor and fragment ions of the most abundant mAb1 heavy chain-derived peptide. (C) Visualization of ion mobility (1/*K*_0_) and *m*/*z* at 38.7 min. (D) Focused visualization of the most abundant mAb1 heavy chain–derived peptide in the ion mobility–*m*/*z* dimension at 38.7 min. (E) Comparison of the total ion chromatograms of mAb1 HCCF and mAb1 PAP. A–D: 30 SPD; E: 15 SPD.

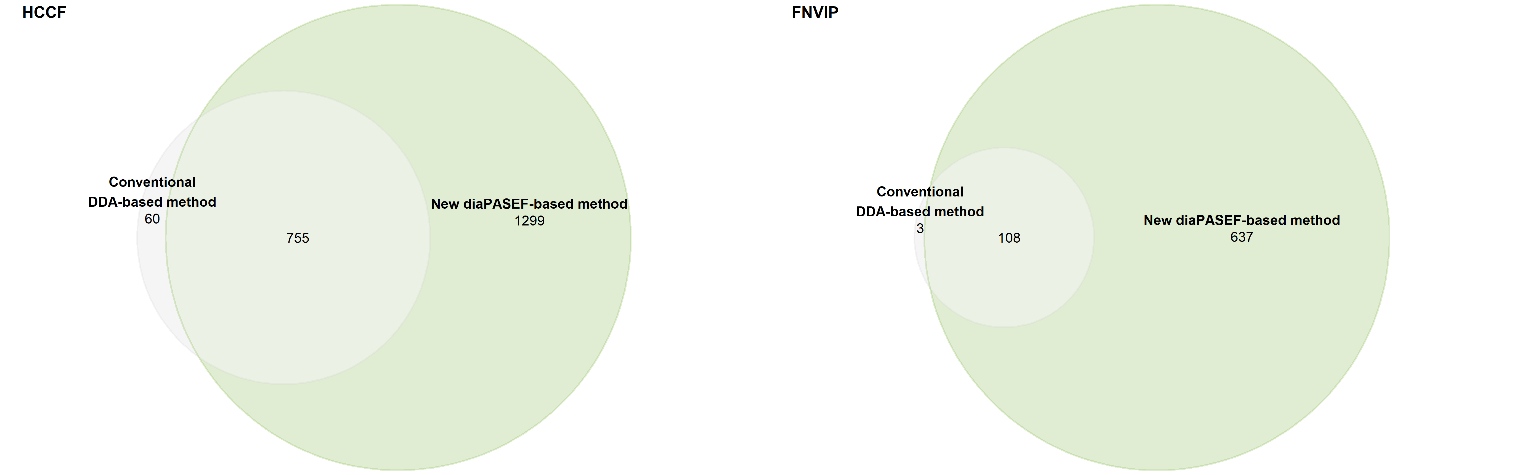

**Figure S8.** Comparison of conventional HCP analysis and new diaPASEF-based HCP analysis. Conventional DDA-based method employs a three-hour gradient on a microflow LC and leverages conventional DDA on Q Exactive HF-X. This method requires 1 mg of sample for processing. mAb3 HCCF and mAb3 filtered neutralized virus inactivation pool (FNVIP) were analysed (N = 3). The numbers of protein groups that were consistently identified in all replicates are shown in the Euler diagrams. Median coefficient of variation in the HCCF data was 21 % by the conventional method and 11 % by the new method while median coefficient of variation in the FNVIP data was 26 % by the conventional method and 19 % by the new method. Missingness in the HCCF data was 16 % by the conventional method and 3 % by the new method while missingness in the FNVIP data was 12 % by the conventional method and 3 % by the new method. Global peptide and global protein FDRs < 1 %.

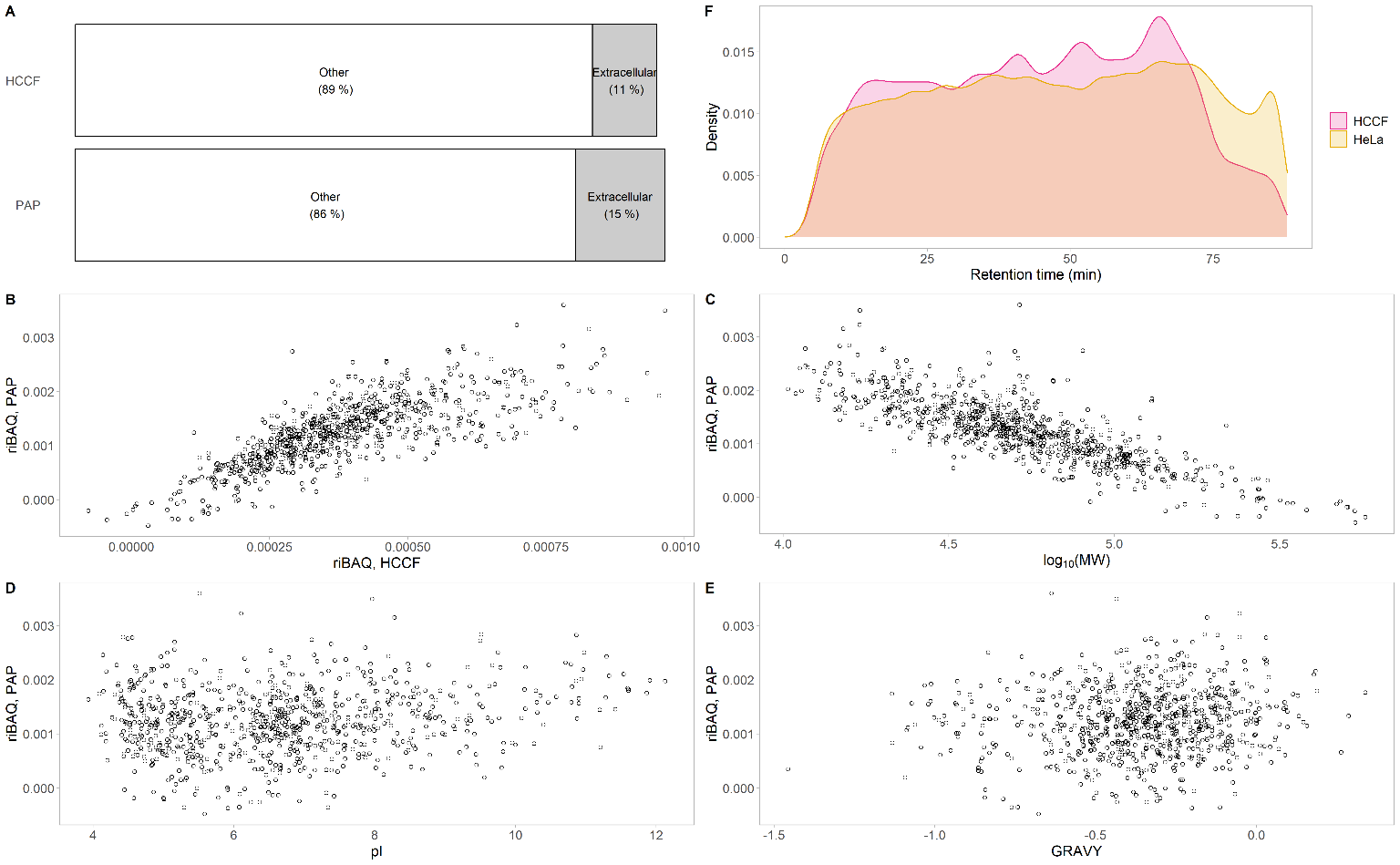

**Figure S9.** Properties of HCPs. (A) Subcellular localization of HCPs quantified in mAb1 HCCF or mAb1 PAP. If HCPs’ human orthologs are annotated with GO:0005615 extracellular space, these HCPs were considered “Extracellular” proteins or otherwise considered “Other”. The percentages were calculated based on abundance (riBAQ). Correlation between abundance in PAP and abundance in HCCF (Spearman’s ρ = 0.79, B), molecular weight (Spearman’s ρ = -0.81, C), isoelectric point (Spearman’s ρ = 0.14, D), or hydrophobicity index (Spearman’s ρ = 0.10, E). (F) Comparison of retention time distribution between HCP (HCCF)-derived peptides and HeLa protein digest.

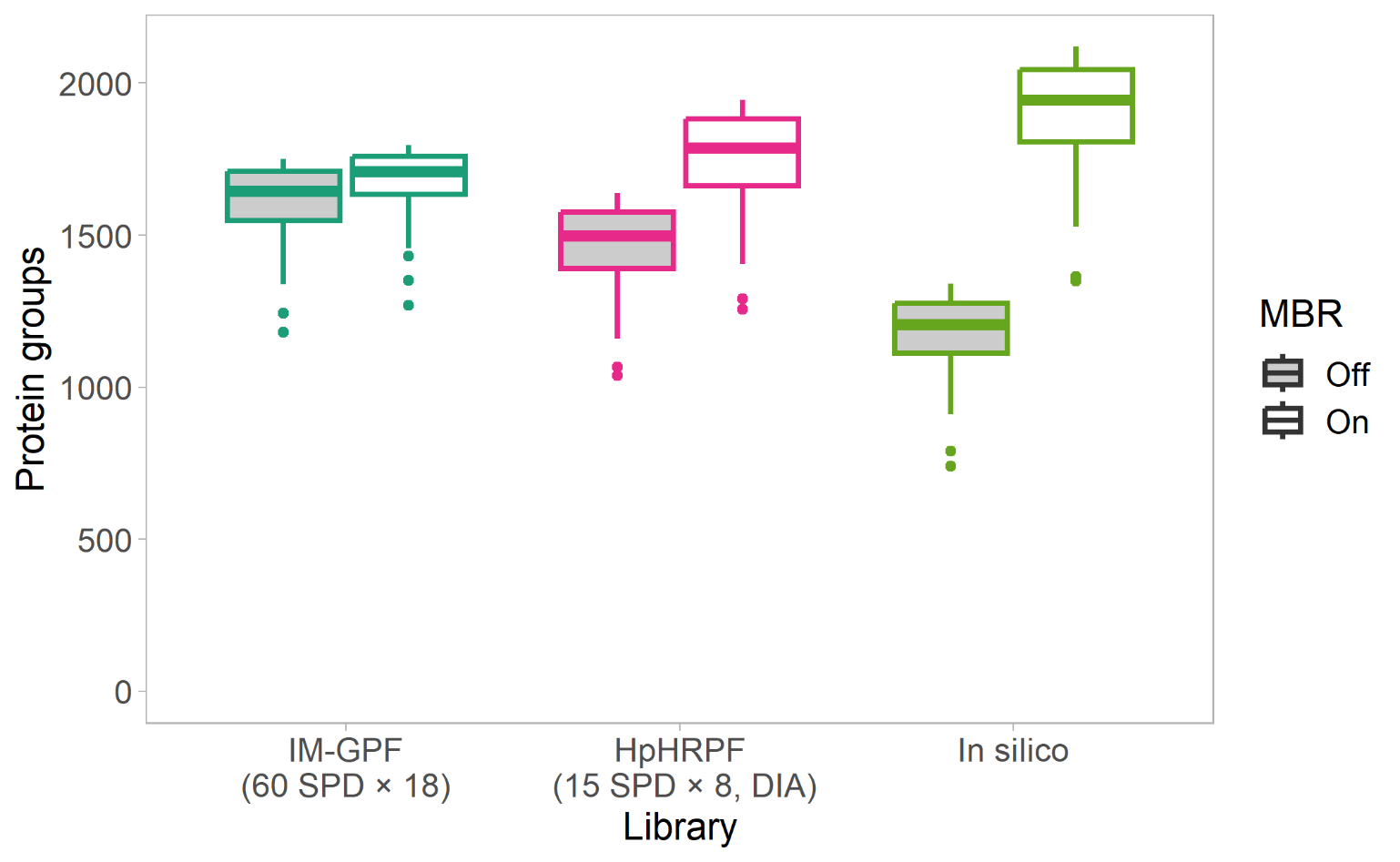

**Figure S10.** Comparison of data processing strategies for high-throughput HCP analysis.
